# Pervasive positive and negative feedback regulation of insulin-like signaling in *Caenorhabditis elegans*

**DOI:** 10.1101/341685

**Authors:** Colin S. Maxwell, Rebecca E. W. Kaplan, Nicole Kurhanewicz Codd, L. Ryan Baugh

## Abstract

The *C. elegans* insulin-like signaling network supports homeostasis and developmental plasticity. The genome encodes 40 insulin-like peptides and one receptor. Feedback regulation has been reported, but the extent of feedback and its effect on signaling dynamics during a state transition has not been determined. We measured mRNA expression for each insulin-like peptide, the receptor *daf-2*, components of the PI3K pathway, and its transcriptional effectors *daf-16*/FoxO and *skn-1*/Nrf at high temporal resolution during transition from a starved, quiescent state to a fed, growing state in wild type and mutants affecting *daf-2*/InsR and *daf-16*/FoxO. We also analyzed the effect of temperature on insulin-like gene expression. We found that numerous PI3K pathway components and insulin-like peptides are affected by signaling activity, revealing pervasive positive and negative feedback regulation. Reporter gene analysis demonstrated that the *daf-2*/InsR agonist *daf-28* positively regulates its own expression and that other agonists cross-regulate *daf-28* transcription through feedback. Our results show that feedback regulation of insulin-like signaling is widespread, suggesting a critical role of feedback in signaling dynamics in this endocrine network and likely others.

## Introduction

Insulin-like signaling maintains homeostasis by responding to fluctuations in nutrient availability and altering gene expression. Work in *C. elegans* has shown that insulin-like signaling also allows developmental plasticity. For example, insulin-like signaling regulates whether larvae become reproductive or arrest as dauer larvae, a developmental diapause that occurs in unfavorable conditions (Hu, 2007). Insulin-like signaling also contributes to continuous variations in phenotype, for example in regulation of aging and growth rate (Murphy and Hu, 2013). However, it is unclear how signaling dynamics are regulated such that the pathway can maintain a phenotypic steady-state (homeostasis) or promote developmental plasticity, depending on conditions.

Insulin-like signaling is regulated by feedback in diverse animals. Pancreatic β-cell-specific insulin receptor-knockout mice are poor at glucose sensing, have a diminished insulin secretory response, and tend to develop age-dependent diabetes (Otani *et al*, 2004). In addition, the full effect of glucose on pancreatic β-cells grown *in vitro* requires the insulin receptor (Assmann *et al*, 2009). FoxO transcription factors, effectors of insulin signaling, activate transcription of insulin receptors in Drosophila and mammalian cells (Puig and Tjian, 2005), suggesting a relatively direct, cell-autonomous mechanism for feedback regulation. However, evidence for such direct feedback regulation has not been found in *C. elegans* (Kimura *et al*, 2011).

Insulin-like signaling regulates the expression of insulin-like peptides in *C. elegans*, suggesting a relatively indirect, cell-nonautonomous mechanism for feedback regulation. The *C. elegans* genome encodes a family of 40 insulin-like peptides that can function as either agonists or antagonists of the sole insulin-like receptor *daf-2* (Pierce *et al*, 2001). Systematic analyses of insulin-like peptide expression and function suggest substantial functional specificity rather than global redundancy (Fernandes de Abreu *et al*, 2014; Ritter *et al*, 2013). *daf-2*/InsR signals through a conserved phosphoinositide 3-kinase (PI3K) pathway to antagonize the FoxO transcription factor *daf-16* (Fig. 1A; Murphy and Hu, 2013)*. daf-16*/FoxO represses transcription of the *daf-2* agonist *ins-7*, creating positive feedback (Murphy *et al*, 2003). This positive feedback results in “FoxO-to-FoxO” signaling, which has been proposed to coordinate the physiological state of different tissues in the animal (Alic *et al*, 2014; Murphy *et al*, 2007; Zhang *et al*, 2013). *daf-16* also activates transcription of the *daf-2* antagonist *ins-18*, again producing positive feedback (Matsunaga *et al*, 2012a; Murphy *et al*, 2003). Insulin-like peptide function has been reported to affect insulin-like peptide expression (Fernandes de Abreu *et al*, 2014; Ritter *et al*, 2013), consistent with feedback regulation. To the best of our knowledge, negative feedback regulation has not been reported, despite the fact that homeostasis generally relies on it (Cannon, 1929). Furthermore, the extent of feedback regulation, and whether it is positive or negative with respect to pathway activity, is unknown.

**Figure 1.**
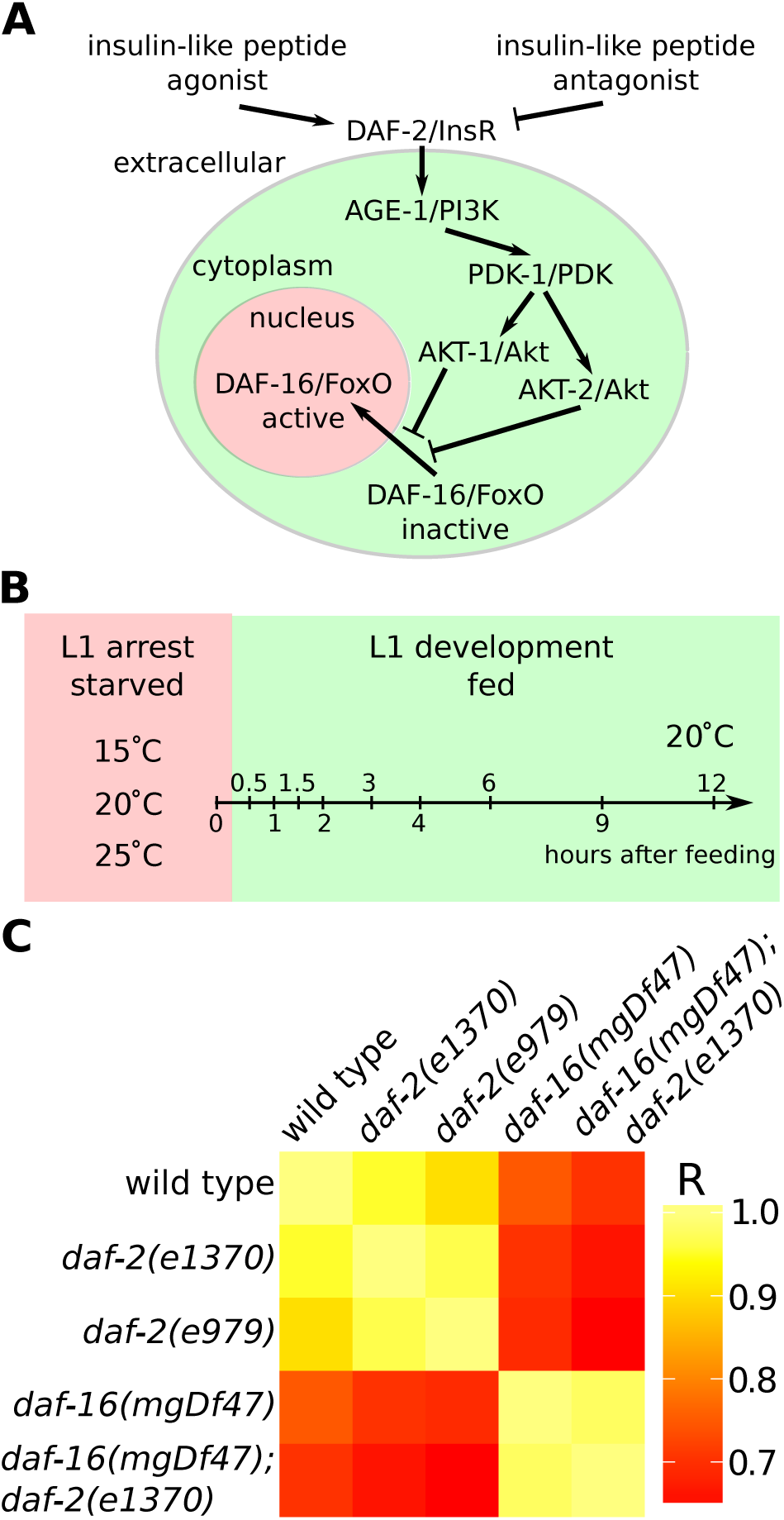
*daf-16*/FoxO is epistatic to *daf-2*/InsR for expression of genes involved in insulin-like signaling. A) A diagram of the C. *elegans* insulin-like signaling pathway. B) A schematic of the experimental design with times and conditions sampled indicated. C) A symmetric matrix of correlation coefficients for pairs of genotypes is presented as a heat map, with scale bar. Examination of individual gene expression patterns confirmed that *daf-16* is epistatic to *daf-2* in each instance without exception.

We sought to determine the extent of feedback regulation in insulin-like signaling in *C. elegans*. *C. elegans* larvae that hatch in the absence of food arrest development in the first larval stage (“L1 arrest” or “L1 diapause”), and insulin-like signaling regulates L1 arrest and development (Baugh, 2013). We performed a genetic analysis of gene expression, measuring expression of all 40 insulin-like peptides as well as components of the PI3K pathway in *daf-2*/InsR and *daf-16*/FoxO mutants, which have perturbed signaling activity. We analyzed larvae in L1 arrest and over time after feeding, as they transition from quiescence to growth. The rationale is that by identifying genes whose expression is affected by insulin-like signaling that themselves affect signaling activity we can infer feedback regulation. We report extensive feedback, both positive and negative, acting relatively directly at the level of the PI3K pathway and also indirectly via regulation of peptide expression. This work suggests that feedback regulation of insulin-like signaling is pervasive and that this feedback functions to stabilize signaling activity during constant conditions while allowing rapid responses to new conditions.

## Results

### *daf-2*/InsR acts through *daf-16*/FoxO to affect gene expression

We used the NanoString nCounter platform to measure expression of genes related to insulin-like signaling in fed and starved L1 larvae at high temporal resolution during the transition between developmental arrest and growth (Malkov *et al*, 2009). Total RNA was prepared from whole worms and hybridized to a codeset containing probes for all 40 insulin-like genes as well as components of the PI3K pathway and *sod-3*, a known DAF-16/FoxO target. In addition to wild type (WT), we analyzed mutations affecting *daf-2*/InsR and *daf-16*/FoxO to ascertain the effects of insulin-like signaling activity on expression. We used the reference allele of *daf-2*, e1370, as well as a stronger allele, e979 (Gems *et al*, 1998). We used a null allele of *daf-16*, mgDf47, as well as a *daf-16(mgDf47); daf-2(e1370)* double mutant to analyze epistasis. Mutations affecting *daf-2* are generally temperature sensitive, and insulin-like signaling responds to temperature. We therefore measured expression during L1 starvation at three different temperatures. We also fed bacteria to starved L1 larvae of each of the five genotypes and measured gene expression over time during recovery from arrest in a highly synchronous population (Fig. 1B). This experimental design enabled us to measure the effects of temperature, nutrient availability, and insulin-like signaling activity on genes related to insulin-like signaling itself during a critical physiological state transition.

*daf-16*/FoxO mediates the effects of *daf-2*/InsR on expression of genes involved in insulin-like signaling. *daf-16* is required for canonical effects of *daf-2*, such as dauer formation and lifespan extension (Hu, 2007; Murphy and Hu, 2013). However, *daf-2* also acts through other effector genes of the PI3K pathway, such as *skn-1*/Nrf (Tullet *et al*, 2008), as well as other signaling pathways, such as RAS (Nanji *et al*, 2005). In addition, genome-wide expression analyses of *daf-16* have mostly been performed in a *daf-2* mutant background (*daf-2* vs. *daf-16*; *daf-2*) without analysis of WT and/or *daf-16* single mutants (Tepper *et al*, 2013), making analysis of epistasis between *daf-2* and *daf-16* with gene expression as a phenotype impossible. Since epistasis was not analyzed, these studies could not determine whether *daf-16* mediated all of the effects of *daf-2* on gene expression or if other effectors made a significant contribution. A correlation matrix between genotypes over all conditions tested indicates that mutating *daf-2* affected expression, with a stronger effect of the e979 allele than e1370, as expected (Fig. 1C). *daf-16* also had a clear effect, and it was epistatic to *daf-2.* That is, the expression profile of the double mutant is similar to that of the *daf-16* single mutant but not *daf-2*. Statistical analysis of individual genes together with examination of expression patterns across genotypes corroborated the results of correlation analysis, failing to identify genes with significant effects of *daf-2* not meditated by *daf-16.* These results show that *daf-2* affects expression of genes involved in insulin-like signaling and that these effects are mediated exclusively by *daf-16*, consistent with feedback regulation.

### *daf-16*/FoxO affects expression of multiple PI3K pathway genes

We analyzed expression of several components of the PI3K pathway, as well as *daf-2*/InsR and its transcriptional effectors *daf-16*/FoxO and *skn-1*/Nrf (Lin *et al*, 1997; Ogg *et al*, 1997; Tullet *et al*, 2008). The known direct target of DAF-16, *sod-3*/SOD (Oh *et al*, 2006), was up-regulated in *daf-2* mutants and down-regulated in the *daf-16* mutant, with *daf-16* epistatic to *daf-2*, in both starved and fed larvae (Fig. 2, S1 and Table 1). The exemplary behavior of this positive control demonstrates the power of our experimental design. Notably, *daf-16* expression drops to background levels in the *daf-16* deletion mutant (Fig. 2 and S1), as expected. We previously reported that *daf-2* is up-regulated during L1 arrest (Chen and Baugh, 2014). We see here that *daf-2* is actually repressed by *daf-16* (Fig. 2 and S1). Given that *daf-2* is up-regulated during starvation, when *daf-16* is active, this result may be considered paradoxical. Our interpretation is that *daf-2* expression is independently regulated by nutrient availability and *daf-16* in opposing ways, illustrating regulatory complexity of the system. Nonetheless, since DAF-2 antagonizes DAF-16 activity via the PI3K pathway, these results indicate positive feedback between the sole insulin-like receptor and its FoxO transcriptional effector (Table 1). Likewise, *age-1*/PI3K, which transduces *daf-2* signaling activity, was repressed by *daf-16*, also suggesting positive feedback. However, *pdk-1*/PDK, *akt-1*/Akt and *akt-2*/Akt, downstream components of the PI3K pathway, were each activated by *daf-16*, albeit with relatively complex dynamics, suggesting negative feedback. Likewise, *daf-16* expression is reduced in *daf-2* mutants (Fig. 2), where its activity is increased, suggesting it represses its own transcription to produce negative feedback (Table 1). *skn-1*/Nrf expression was also reduced in *daf-2* mutants and increased in *daf-16* mutants, suggesting that insulin-like signaling positively regulates expression of both of its transcriptional effectors. Notably, the effects described here for each gene were consistent for fed and starved larvae (Fig. 2, S1 and Supp. Data File 2). In summary, insulin-like signaling acts through *daf-16*/FoxO to regulate multiple critical components of the pathway itself, consistent with a combination of positive and negative cell-autonomous feedback regulation.

**Figure 2.**
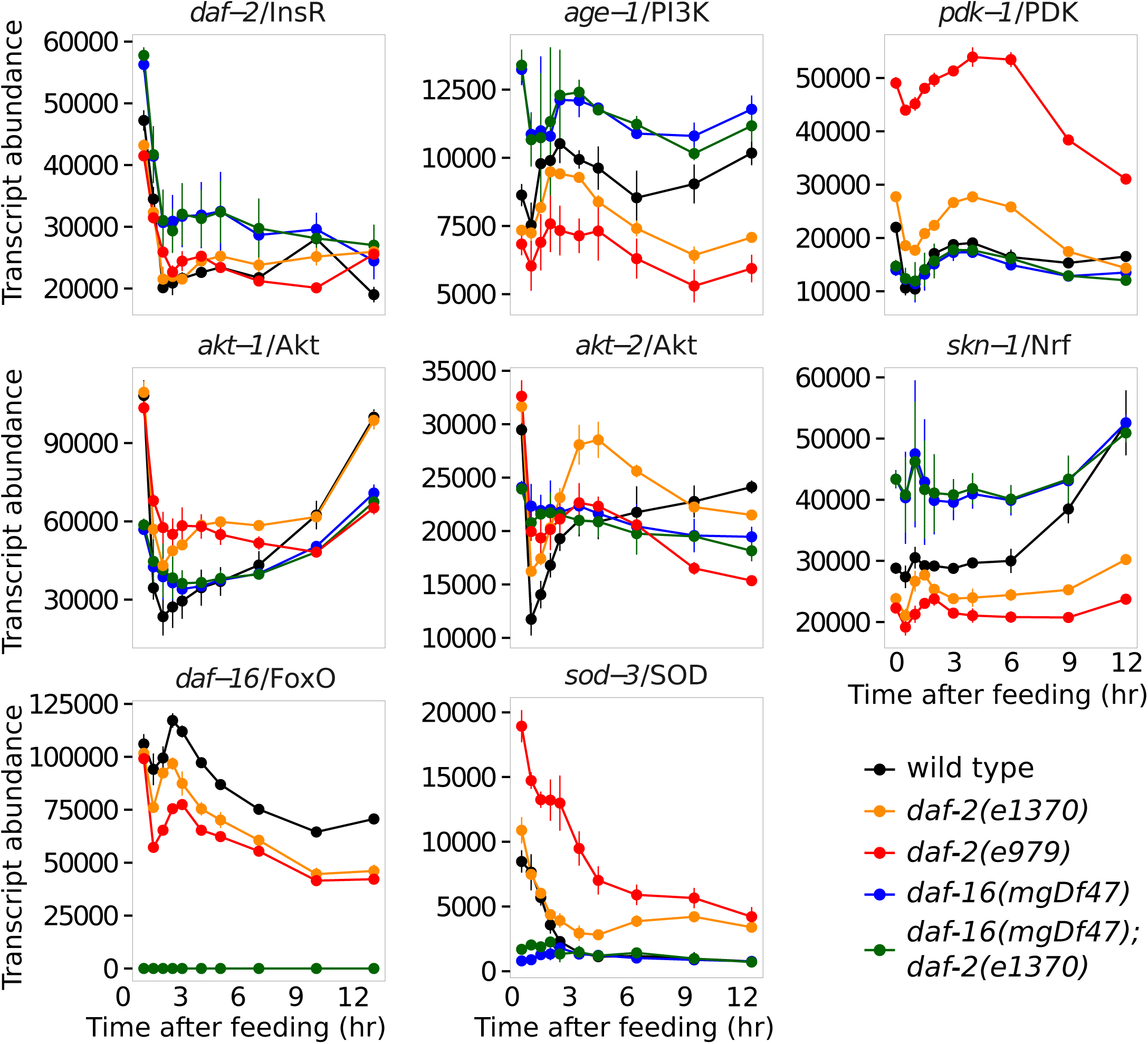
Insulin-like signaling regulates expression of genes comprising the insulin-like signaling pathway. Transcript abundance (arbitrary units) is plotted over time during recovery from L1 starvation by feeding in five different genotypes with various levels of insulin-like signaling activity. In addition to *daf-2*/InsR and components of the PI3K pathway, the transcriptional effectors of signaling, *daf-16*/FoxO and *skn-1*/Nrf, are plotted as well as the known DAF-16 target *sod-3*. Note that *daf-16* expression was not detected in *daf-16* mutants. Each gene plotted was significantly affected by *daf-16* (see Tables 1, S1 and supp. data). Error bars reflect the SEM of two or three biological replicates.

**Table 1.**
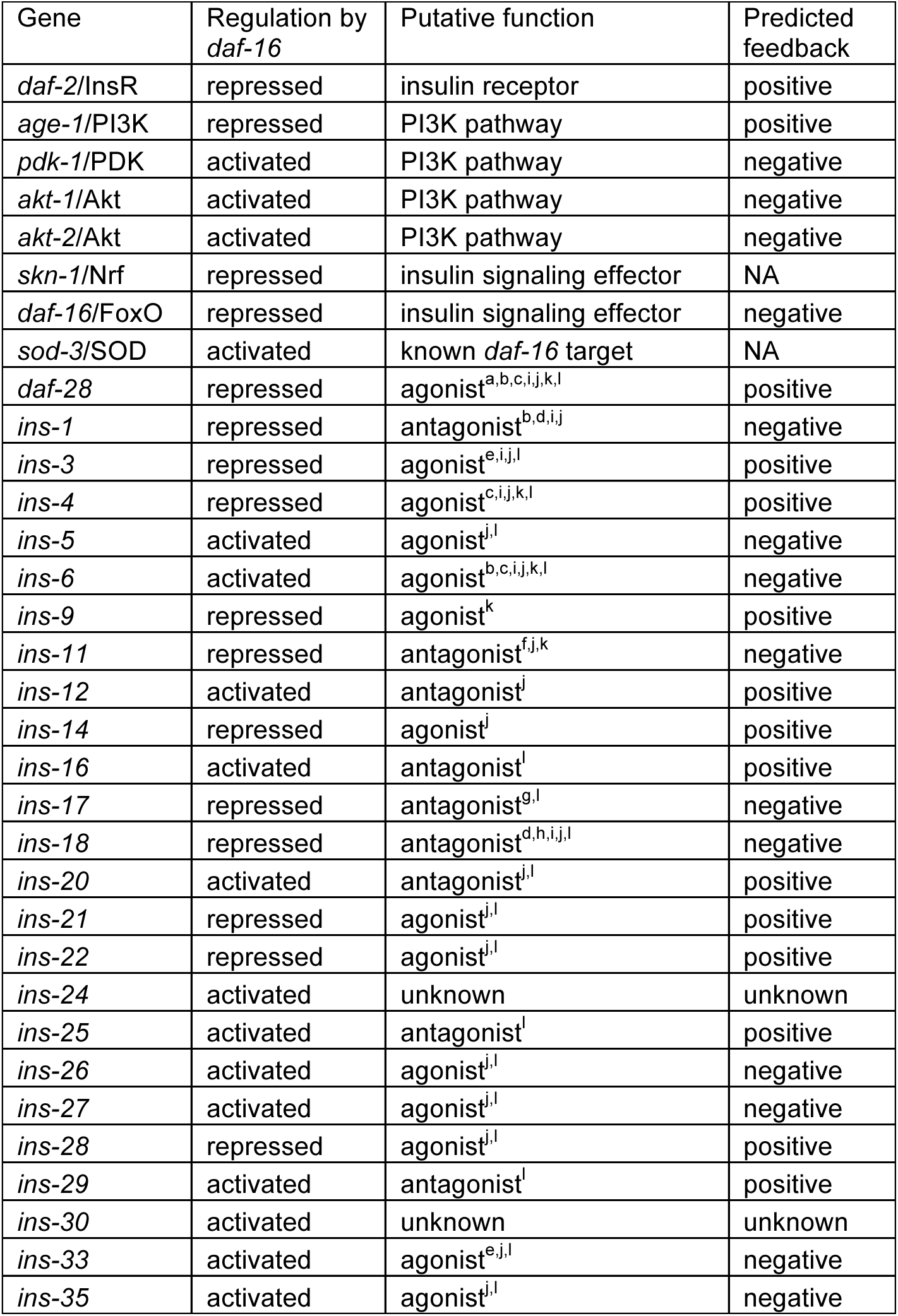
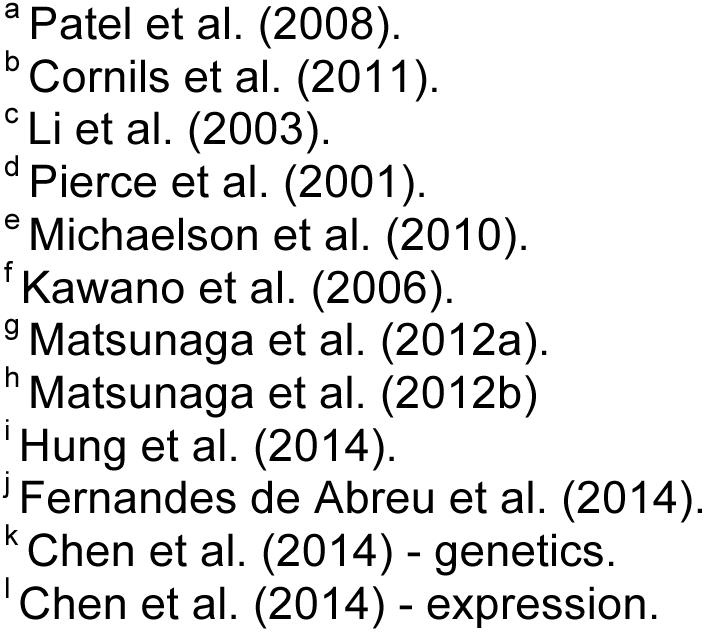
Summary of genes regulated by *daf-16*/FoxO. The gene, whether it is activated or repressed by *daf-16*, putative function of the gene, and whether regulation is predicted to result in positive or negative feedback is presented. Four total tests for regulation were considered (during L1 starvation at three different temperatures and during recovery over time after feeding, comparing *daf-16(mgDf47)* to WT and also *daf-16(mgDf47); daf-2(e1370)* to *daf-2(e1370)*). Results are considered significant if the p-value is below 0.05 in any one test after correction for multiple testing. See supplementary information for complete statistical analysis. Insulin-like peptides are predicted to function as agonists or antagonists of *daf-2*/InsR based on published genetic or expression analysis (Chen et al. 2014 is cited separately for results based on genetic or expression analysis; all other citations are for genetic analysis), and positive or negative feedback is predicted based on putative function (agonist or antagonist) and whether the gene is positively or negatively regulated by *daf-16*.

### *daf-16*/FoxO affects expression of most insulin-like peptides

Insulin-like genes display complex dynamics in response to different levels of insulin-like signaling activity. Our codeset contained probes for all 40 insulin-like genes, and we reliably detected expression for 28 of them. Similar to what we saw with components of the PI3K pathway (Fig. 2, S1 and Table 1), *daf-16* appears to function as an activator in some cases and a repressor in others (Fig. 3, S2 and Table 1), but its function with respect to each gene affected was again consistent between fed and starved conditions (Fig. 3, S2 and Supp. Data File 2). For example, expression of *daf-28*, perhaps the most studied insulin-like peptide in *C. elegans* (Chen and Baugh, 2014; Cornils *et al*, 2011; Fernandes de Abreu *et al*, 2014; Hung *et al*, 2014; Li *et al*, 2003; Patel *et al*, 2008), was up-regulated in *daf-16* mutants and down-regulated in *daf-2* in starved and fed larvae (Fig. 3, S2 and Table 1), suggesting it is repressed by *daf-16.* Remarkably, all but three of the 28 reliably detected insulin-like genes were significantly affected by *daf-16* (Table 1). Mutation of *daf-16* caused up-regulation of twelve insulin-like genes and down-regulation of thirteen, suggesting that *daf-16* directly or indirectly regulates transcription of most insulin-like genes.

**Figure 3.**
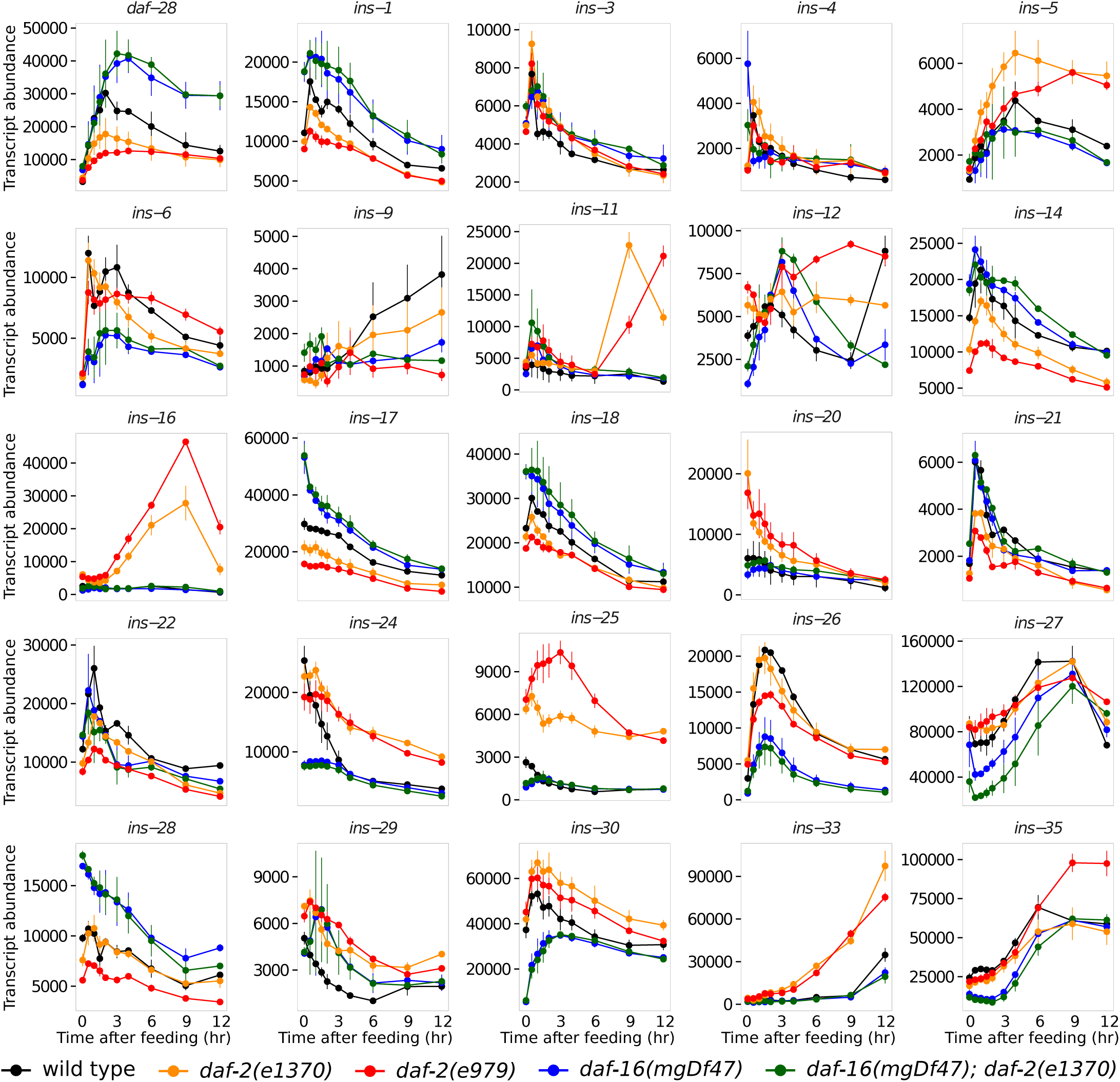
Insulin-like signaling regulates expression of the majority of insulin-like peptides. Transcript abundance (arbitrary units) is plotted over time during recovery from L1 starvation by feeding in five different genotypes with various levels of insulin-like signaling activity. Of 28 reliably detected insulin-like genes, 25 were significantly affected by *daf-16* (all but ins-2, −10 and -*34*; see Tables 1, S1 and supp. data) and are plotted. Error bars reflect the SEM of two or three biological replicates.

Inference of feedback as positive or negative is complicated by the fact that individual insulin-like peptides function as either agonists or antagonists of *daf-2*/InsR (Pierce *et al*, 2001). Biochemical data and structural modeling suggest that function as an agonist or antagonist is a property of the peptide (Matsunaga *et al*, 2018), as opposed to the context in which it is expressed. To infer whether the net effect of feedback regulation is positive or negative with respect to insulin-like signaling activity (*daf-2*/InsR activity), we took into account whether *daf-16* appears to activate or repress the insulin-like gene and whether that gene encodes a putative agonist or antagonist. DAF-2 antagonizes DAF-16 activity, and so *daf-16* repression or activation of an agonist or antagonist, respectively, would hypothetically result in positive feedback. *daf-16* repression or activation of an antagonist or agonist, respectively, would hypothetically result in negative feedback. For example, *daf-28* was originally identified on the basis of its constitutive dauer-formation phenotype. *daf-28* is up-regulated in rich conditions and it promotes dauer bypass (reproductive development), similar to *daf-2*/InsR, consistent with function as an agonist of *daf-2* (Li *et al*, 2003). *daf-16* repression of *daf-28* expression therefore suggests positive feedback in this case (Table 1).

A number of studies have performed genetic analysis of insulin-like peptide function, determining whether individual insulin-like genes have similar or opposite loss-of-function phenotypes to *daf-2*, and thus whether they presumably function as agonists or antagonists, respectively (Chen and Baugh, 2014; Cornils *et al*, 2011; Fernandes de Abreu *et al*, 2014; Hung *et al*, 2014; Kawano *et al*, 2006; Li *et al*, 2003; Matsunaga *et al*, 2012a; Matsunaga *et al*, 2012b; Michaelson *et al*, 2010; Patel *et al*, 2008; Pierce *et al*, 2001). When we previously analyzed expression of insulin-like peptides in starved and fed L1 larvae, we found remarkable concordance between function (agonist or antagonist) and expression (positive or negative effect of food, respectively) (Chen and Baugh, 2014). Out of thirteen insulin-like peptides consistently found to function as putative agonists or antagonists based on genetic analysis, we classified all thirteen the same way based on expression, while classifying eight additional peptides as well. This classification relied on separate time-series analyses of starved and fed larvae (Chen and Baugh, 2014), and inspection of the fed time series here did not reveal discrepancies between the two studies. We therefore included our previous putative functional classifications based on nutrient-dependent expression in Table 1, which tentatively assigns function to all but two of the 25 genes affected by *daf-16*. As explained above, putative agonists repressed by *daf-16*, like *daf-28*, hypothetically result in positive feedback, since *daf-2* signaling antagonizes *daf-16*. We identified seven genes like this in addition to *daf-28.* Conversely, activation of a putative antagonist should also produce positive feedback, which we infer in five cases, while activation of an agonist should produce negative feedback, which we infer in six cases. Finally, repression of a putative antagonist should produce negative feedback, which we infer in four cases. In summary, activation and repression of putative agonists and antagonists by *daf-16* is common, with positive and negative feedback hypothetically resulting from each different regulatory combination in multiple instances.

### Temperature affects insulin-like gene expression

We analyzed expression of insulin-like genes at 15, 20 and 25°C during L1 starvation. *daf-2* mutants are generally temperature-sensitive (Gems *et al*, 1998), *daf-16* is localized to the nucleus at high temperatures (Henderson and Johnson, 2001), and *daf-2* mutants are heat-resistant (Munoz and Riddle, 2003). These observations suggest that insulin-like signaling responds to temperature. We hypothesized that temperature sensitivity results from temperature-dependent regulation of insulin-like peptide expression. Consistent with *daf-16* being active at elevated temperature, expression of its direct target *sod-3* was positively affected by temperature (Fig. S2 and Table S1). In support of our hypothesis, temperature affected mRNA expression of 21 out of 28 reliably detected insulin-like genes (Fig. S2 and Table S1). *daf-28* expression was lower at higher temperatures, consistent with its role in promoting dauer bypass (Li *et al*, 2003), and confirmed in a recent publication (O’Donnell *et al*, 2018). Expression of twelve insulin-like genes was lower at higher temperatures and nine were expressed higher at higher temperatures. However, there is no apparent correlation between putative function as agonist or antagonist and positive or negative regulation in response to higher temperature. Notably, although most insulin-like genes displayed significant temperature-dependent expression, the effect of temperature on expression was minor compared to nutrient availability.

### Feedback mediates cross-regulation among insulin-like genes

Reporter gene analysis validated the effect of *daf-16*/FoxO on *daf-28* expression. We previously used quantitative RT-PCR to validate the nCounter approach to measuring insulin-like gene expression in *C. elegans* (Baugh *et al*, 2011), and we used transcriptional reporter genes to confirm positive regulation of several putative agonists in fed larvae, including *daf-28* (Chen and Baugh, 2014). A P*daf-28*::GFP transcriptional reporter gene again confirmed up-regulation in response to feeding (Fig 4A). Expression was evident but faint in anterior neurons and posterior intestine of starved L1 larvae, and it was brighter after being fed for 6 hr. Quantification of whole-animal fluorescence with the COPAS BioSorter provided robust statistical support for qualitative observations (Fig. 4B). Note that the statistics for this analysis were performed on the means of individual biological replicates, as opposed to each individual in a replicate. Thus, statistical significance is due to reproducibility despite relatively small effect sizes. Critically, expression appeared elevated in *daf-16* mutants compared to WT, in both starved and fed larvae (Fig. 4A). However, we did not observe a difference in the anatomical expression pattern in *daf-16* compared to WT. Quantification showed that the effect of *daf-16* is statistically significant (Fig. 4B). Notably, the effect of food was larger than that of *daf-16*, as expected based on nCounter results (Fig. 3). In addition, the effects of food and *daf-16* are independent, suggesting that up-regulation of *daf-28* in response to feeding is not simply due to inhibition of *daf-16* leading to de-repression of *daf-28*. These results support the conclusion that *daf-16* represses *daf-28* transcription, consistent with feedback regulation.

**Figure 4.**
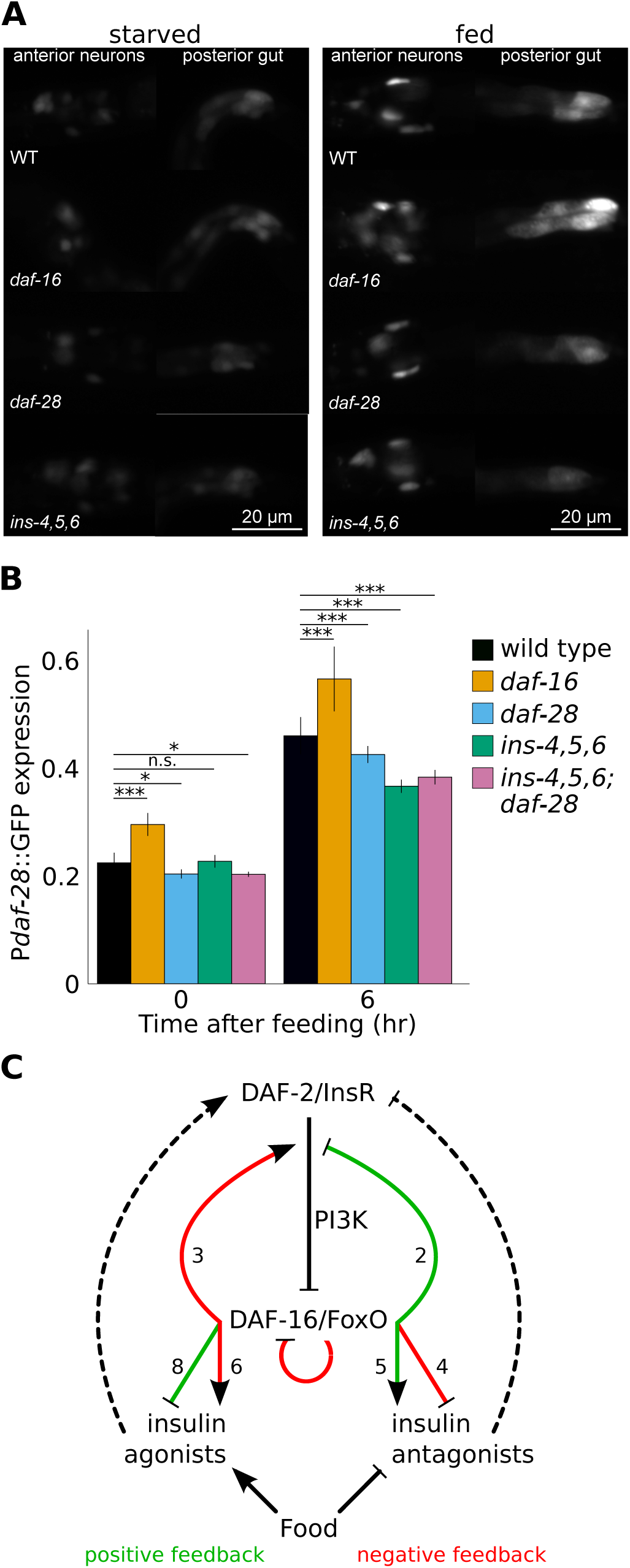
Insulin-like peptide function affects expression of insulin-like peptides. A) Representative images of a P*daf-28*::GFP transcriptional reporter gene are presented for WT, *daf-16(mu86)*, *daf-28(tm2308)* and *ins-4, 5, 6(hpDf761)* in starved and fed (6 hr) L1 larvae, as indicated. Images were cropped and expression in anterior neurons and posterior gut is shown. B) Quantitative analysis of P*daf-28*::GFP expression using the COPAS BioSorter is presented. The grand average and standard deviation of three to seven biological replicates is plotted for starved (0 hr) and fed (6 hr) L1 larvae. *p<0.05, ***p<0.001 (unpaired t-test on replicate means, n=3 to 7). C) Pervasive feedback regulation of insulin-like signaling. *daf-2*/InsR antagonizes *daf-16*/FoxO via the PI3K pathway. *daf-16* activates expression of three pathway components to produce negative feedback, and it represses expression of two components to produce positive feedback. *daf-16* also appears to activate its own expression, producing positive feedback. *Daf-16* represses expression of eight putative insulin-like peptide agonists of *daf-2* and activates five putative antagonists, producing positive feedback in each case. *daf-16* also activates six putative agonists and represses four putative antagonists, producing negative feedback. Food activates expression of agonists and represses expression of antagonists, independent of *daf-16* and insulin-like signaling. Dashed arrows reflect putative function of insulin-like peptides as agonists or antagonists of *daf-2*, reflecting cell-nonautonomous effects of *daf-16*. Inferred positive feedback is depicted in green and negative feedback in red. Numbers next to colored arrows indicate the number of genes represented by each arrow.

Widespread feedback regulation of insulin-like signaling via transcriptional control of insulin-like peptides suggests that activity of individual insulin-like genes should affect expression of themselves and others. We analyzed expression of P*daf-28*::GFP in insulin mutants to test this hypothesis. P*daf-28*::GFP transgene expression was significantly reduced in a *daf-28* mutant (Fig. 4A,B). This result suggests that positive feedback mediated by *daf-16* repression of *daf-28*, *daf-28* agonism of *daf-2*/InsR, and *daf-2* inhibition of *daf-16* results in *daf-28* effectively promoting its own expression. *daf-28*, *ins-4* and *ins-6* coordinately regulate dauer entry and exit (Cornils *et al*, 2011), and they redundantly promote L1 development in response to feeding (Chen and Baugh, 2014). *ins-4, 5* and *6* are in a chromosomal cluster, so we analyzed a deletion allele that removes all three (Hung *et al*, 2014). P*daf-28*::GFP expression was significantly reduced in fed larvae of the *ins-4, 5, 6* mutant compared to WT (Fig. 4A,B). This result suggests that feedback regulation results in cross-regulation among insulin-like peptides such that the function of one peptide affects the expression of others. Compound mutants affecting *ins-4, 5, 6* and *daf-28* grow slowly as fed L1 larvae and display starvation resistance during L1 arrest (Chen and Baugh, 2014), and P*daf-28*::GFP expression was also reduced consistent with these phenotypes (Fig. 4B). In summary, reporter gene analysis suggests physiological significance of feedback regulation, consistent with function of individual insulin-like peptides affecting expression of others.

## Discussion

We determined the extent of feedback regulation of insulin-like signaling in *C. elegans* in starved and fed L1 larvae. We show that mRNA expression of nearly all detectable insulin-like genes is affected by insulin-like signaling activity, revealing pervasive feedback regulation. We also show that several components of the PI3K pathway, including *daf-2*/InsR and *daf-16*/FoxO, are affected by signaling activity. Together these results suggest that feedback occurs inter- and intra-cellularly (Fig. 4C). Furthermore, we show that feedback is positive and negative at both levels of regulation. Finally, we demonstrate that feedback regulation results in auto- and cross-regulation of insulin-like gene expression.

We detected substantially more regulation of insulin-like genes by *daf-16*/FoxO than previously reported in genome-wide expression analyses. We also detected extensive effects of temperature on insulin-like gene expression. In contrast to other expression analyses, our analysis employed highly synchronous populations of larvae, improving sensitivity. Sensitivity was also likely improved by focusing on proximal effects of nutrient availability, which has robust effects on insulin-like signaling. In addition, the nCounter assay conditions used are optimized for sensitivity and precision (Baugh *et al*, 2011), improving power to detect differential expression. We also analyzed the effects of *daf-16* mutation in a WT background as well as a *daf-2* mutant background, in fed and starved larvae, producing four independent opportunities to detect an effect of *daf-16.* Finally, we sampled extensively, not only with biological replicates, but also with three different temperatures during L1 arrest as well as nine time points after feeding. Taken together, these features likely explain why we detected such extensive effects.

Other nutrient-dependent pathways also regulate expression of insulin-like genes and PI3K pathway components. That is, insulin-like signaling does not account for all of the observed effects of nutrient availability on gene expression (Fig. 4C). For example, we show that *daf-28* expression is up-regulated in response to feeding and that it is repressed by *daf-16*/FoxO. Since DAF-16 is nuclear and active during starvation and is excluded from the nucleus in response to feeding (Henderson and Johnson, 2001), it is conceivable that up-regulation of *daf-28* in response to feeding is due to inactivation of DAF-16 and de-repression of *daf-28.* However, this model predicts that *daf-28* expression should be equivalent in starved and fed *daf-16* mutant larvae, but it is not. To the contrary, induction of *daf-28* in fed larvae occurs with similar magnitude in each genotype tested. This was true with mRNA expression analysis by nCounter as well as transcriptional reporter gene analysis. Despite numerous examples of ***daf-2*** and ***daf-16*** affecting expression, the effects of nutrient availability are generally evident in all genotypes, indicating the influence of other nutrient-dependent pathways (Fig. 4C).

We provide evidence that *daf-16*/FoxO activity leads to activation and repression of genes involved in insulin-like signaling. Both modes of regulation were observed for putative *daf-2*/InsR agonists and antagonists, supporting the conclusion agonists and antagonists both contribute to positive and negative feedback regulation. However, we used genetic and not biochemical analysis, so we do not know if DAF-16 regulation is direct or indirect. DAF-16 is thought to function primarily as an activator (Riedel *et al*, 2013; Schuster *et al*, 2010), with repression (“class II” targets) occurring indirectly via its antagonism of the transcriptional activator PQM-1 (Tepper *et al*, 2013). However, a role of *pqm-1* in L1 arrest and recovery has not been investigated. Nonetheless, *akt-1*/Akt, *akt-2*/Akt, *skn-1*/Nrf and *daf-16*/FoxO were each included on a list of 65 high-confidence direct DAF-16 targets (Schuster *et al*, 2010). We found each of these to be regulated by *daf-16*, with *skn-1* and *daf-16* being repressed, consistent with direct repression independent of PQM-1. Mechanistic details aside, this work reveals extensive positive and negative feedback regulation of insulin-like signaling.

Insulin-like peptide function regulates expression of insulin-like genes. We used reporter gene analysis to show that function of *daf-28*, a *daf-2* agonist repressed by *daf-16*, affects its own transcription. Furthermore, we showed that function of other agonists cross-regulate *daf-28* transcription. These results are consistent with reports of insulin-like peptides affecting expression of insulin-like genes (Fernandes de Abreu *et al*, 2014; Ritter *et al*, 2013), though in this case we demonstrate an intermediary effect of *daf-16*/FoxO. Given that we found most insulin-like genes to be regulated by insulin-like signaling, cross regulation among insulin-like peptides is likely common.

We believe the physiological significance of feedback regulation is to stabilize signaling activity in variable environments. Negative feedback supports homeostasis, returning the system to a stable steady state (Cannon, 1929). In contrast, positive feedback supports rapid responses and switch-like behavior (Ingolia and Murray, 2007). We speculate that by combining negative and positive feedback, the insulin-like signaling system is able to maintain homeostasis at different set points of signaling activity. That is, in constant conditions negative feedback stabilizes signaling activity, but when conditions change (e.g., differences in nutrient availability) positive feedback allows signaling activity to respond rapidly and negative feedback helps it settle to a new steady state rather than displaying runaway dynamics. In addition, signaling occurs in the context of a multicellular animal, with tissues and organs that presumably vary in their energetic and metabolic demands. Consequently, FoxO-to-FoxO signaling resulting from feedback may be relatively positive or negative in different anatomical regions, governed by the peptides involved, serving to coordinate the animal’s physiology appropriately (Kaplan and Baugh, 2016; McMillen *et al*, 2002). In any case, the extent of feedback suggests that it is a very important means of regulation. We imagine that insulin-like signaling in other animals and other endocrine signaling systems are also rife with feedback, and that it is critical to system dynamics.

## Materials and Methods

### Nematode culture and sample collection

The following *C. elegans* strains were used for gene expression analysis on the NanoString nCounter platform: N2 (wild type), PS5150 (*daf-16(mgDf47)*), CB1370 (*daf-2(e1370)*), DR1942 (*daf-2(e979)*), GR1309 (*daf-16(mgDf47); daf-2(e1370)*). Strains were maintained on NGM agar plates with *E. coli* OP50 as food at 15°C (DR1942) or 20°C (all others). Liquid culture was used to obtain sufficiently large populations for time-series analysis with microgram-quantities of total RNA. Larvae were washed from clean, starved plates with S-complete and used to inoculate liquid cultures (Lewis, 1995). A single 6 cm plate was typically used, except with CB1370 and DR1942, for which two and three plates were used, respectively. Liquid cultures were comprised of S-complete and 40 mg/ml *E. coli* HB101. These cultures were incubated at 180 rpm and 15°C for four days (with the exception of DR1942, which was incubated for five days), and eggs were prepared by standard hypochlorite treatment, yielding in excess of 100,000 eggs each. These eggs were used to set up another liquid culture again consisting of S-complete and 40 mg/ml HB101 but with a defined density of 5,000 eggs/ml. These cultures were incubated at 180 rpm and 15°C for five days (N2, PS5150 and GR1309), six days (CB1370) or seven days (DR1942), and eggs were prepared by hypochlorite treatment with yields in excess of one million eggs per culture. These eggs were cultured in S-complete without food at a density of 5,000 eggs/ml at 180 rpm so they hatch and enter L1 arrest. For starved samples at 20°C and 25°C, they were cultured for 24 hr and collected, and for 15°C they were cultured for 48 hr. Fed samples were cultured for 24 hr at 20°C, and then 25 mg/ml HB101 was added to initiate recovery by feeding. Fed samples were collected at the time points indicated. Upon collection, larvae were quickly pelleted by spinning at 3,000 rpm for 10 sec, washed with S-basal and spun three times, transferred by Pasteur pipet to a 1.5 ml plastic tube in 100 µl or less, and flash frozen in liquid nitrogen. Samples were collected in at least two but typically three independent biological replicates where the entire culture and collection process was repeated.

### RNA preparation and hybridization

Total RNA was prepared using 1 ml TRIzol (Invitrogen) according to the manufacturer’s instructions. 3 µg total RNA was used for hybridization by NanoString, Inc (Seattle, WA USA), as described (Chen and Baugh, 2014). The codeset used included the same probes for all insulin-like genes as in Chen, 2014 with the exception of *ins-13*, which was replaced here. The codeset also included probes for additional genes not included in Chen, 2014 (for a complete list of genes targeted see Supplementary Data File 1) as well as standard positive and negative control probes.

### Data analysis

nCounter results were normalized in a two-step procedure. First, counts for positive control probes (for which transcripts were spiked into the hybridization at known copy numbers) were used to normalize the total number of counts across all samples. Second, the total number of counts for all targeted genes except *daf-16* (the deletion mutant used did not produce signal above background) was normalized across all samples. Insulin genes with a normalized count of less than 5,000 were excluded from further analysis because they displayed a cross-hybridization pattern indicating that they were not reliably detected. The complete normalized data set is available in Supplementary Data File 1.

Statistical analysis was used to assess the effects of *daf-16* (in fed and starved samples) and temperature (starved samples only). For the effect of *daf-16* in fed samples, two tests were used: a non-parametric ANCOVA with the null hypothesis that loess lines connecting the points of the *daf-16* single mutant (or the *daf-16; daf-2* double mutant) and wild type (or *daf-2(e1370)*) are overlapping. This test was implemented using the R package “sm” (Bowman and Azzalini, 1997). For the effect of *daf-16* in starved samples, two tests were used: a bootstrap test was used with the null hypothesis that the *daf-16* single mutant (or the *daf-16; daf-2* double mutant) has the same mean expression level as wild type (or *daf-2(e1370)*) for all temperatures. The effect size of genotype is calculated within each temperature, so it controls for temperature. 10,000 permutations of genotype were calculated to get the p-value. For the effect of temperature during starvation, a chi-squared goodness of fit test was used to ask whether temperature explained additional variance in gene expression after controlling for genotype. Benjamani-Hochberg was used to calculate the ‘q-value’ (Benjamini and Hochberg, 1995), and these q-values were used to identify genes affected by *daf-16* or temperature at a false-discovery rate of 5%. The complete results of statistical analysis is available in Supplementary Data File 2.

### Reporter gene analysis

The mgIs40 [P*daf-28*::GFP] reporter (Li *et al*, 2003) was analyzed using the following genetic backgrounds: wild type (N2), *daf-16(mu86)*, *daf-28(tm2308)* and *ins-4, 5, 6(hpDf761)*. Strains were maintained on NGM agar plates with *E. coli* OP50 as food at 20°C. Eggs were prepared by standard hypochlorite treatment. These eggs were used to set up a liquid culture consisting of S-basal without ethanol or cholesterol with a defined density of 1,000 eggs/ml. After 18 hours to allow for hatching, *E. coli* HB101 was added at 25 mg/ml to the fed samples. 6 hours post food addition, the samples were washed three times with 10 ml S-basal and then run through the COPAS BioSorter measuring GFP fluorescence. Analysis of the COPAS data was performed in R. Tukey fences were used to remove outliers. Data points were also removed if they were determined to be debris by size or lack of fluorescent signal. This cleanup left a total of almost 165,000 data points. Fluorescence data was normalized by worm density. The Bartlett test of homogeneity of variances rejected the null hypothesis that the samples had equal variance. Therefore, unpaired t-tests with unequal variance were used to determine the significance of condition and genotype on mean normalized fluorescence. There were three biological replicates for the insulin-like peptide mutants and seven biological replicates for wild type and *daf-16* mutants.

For imaging, the samples were prepared in the same way then paralyzed with 3.75 mM sodium azide and placed on an agarose pad on a microscope slide. Images were taken on a compound fluorescent microscope.

## Data Availability

The complete normalized data set is available in Supplementary Data File 1. Complete results of statistical analysis is available in Supplementary Data File 2. Raw data and strains used here are available upon request.

## Acknowledgements

We would like to thank NanoString, Inc. for performing nCounter hybridizations for us. Some strains were provided by the CGC, which is funded by NIH Office of Research Infrastructure Programs (P40 OD010440). The National Science Foundation (IOS-1120206) and the National Institutes of Health (R01GM117408) funded this work.

## Author Contributions

LRB conceived of the study and provided funding. LRB, REWK and NKC performed the experiments. CSM and REWK analyzed the data. LRB, CSM and REWK prepared the manuscript.

---

**Figure S1 - complementing Fig. 2.**
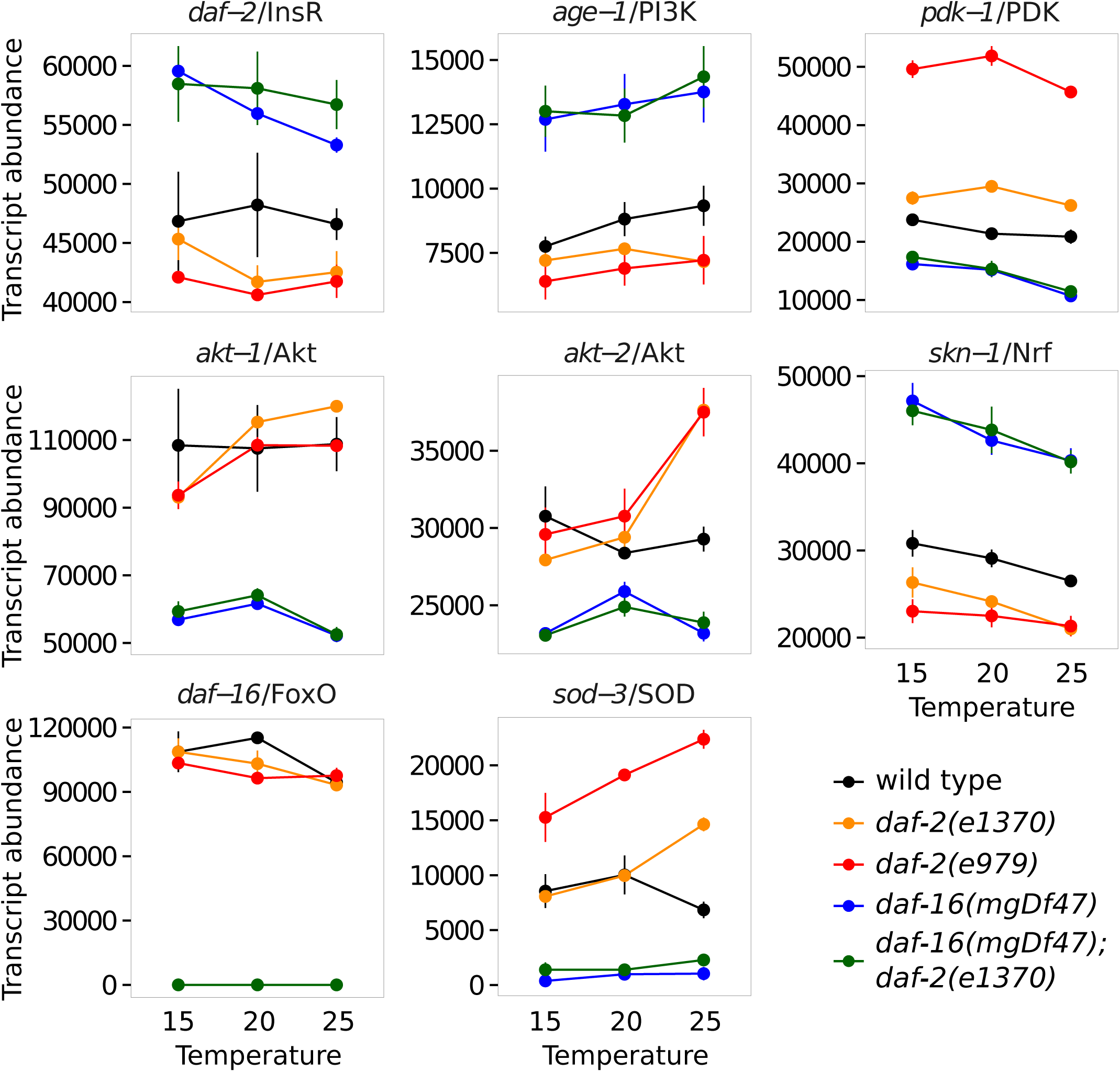
Insulin-like signaling regulates expression of genes comprising the insulin-like signaling pathway during L1 starvation. Transcript abundance (arbitrary units) is plotted during L1 starvation at three different temperatures in five different genotypes with various levels of insulin-like signaling activity. In addition to *daf-2*/InsR and components of the PI3K pathway, the transcriptional effectors of signaling, *daf-16*/FoxO and *skn-1*/Nrf, are plotted as well as the known DAF-16 target *sod-3*. Note that *daf-16* expression was not detected in *daf-16* mutants. Each gene plotted was significantly affected by *daf-16* (see Tables 1, S1 and supp. data). Error bars reflect the SEM of two or three biological replicates.

**Figure S2 - complementing Fig. 3.**
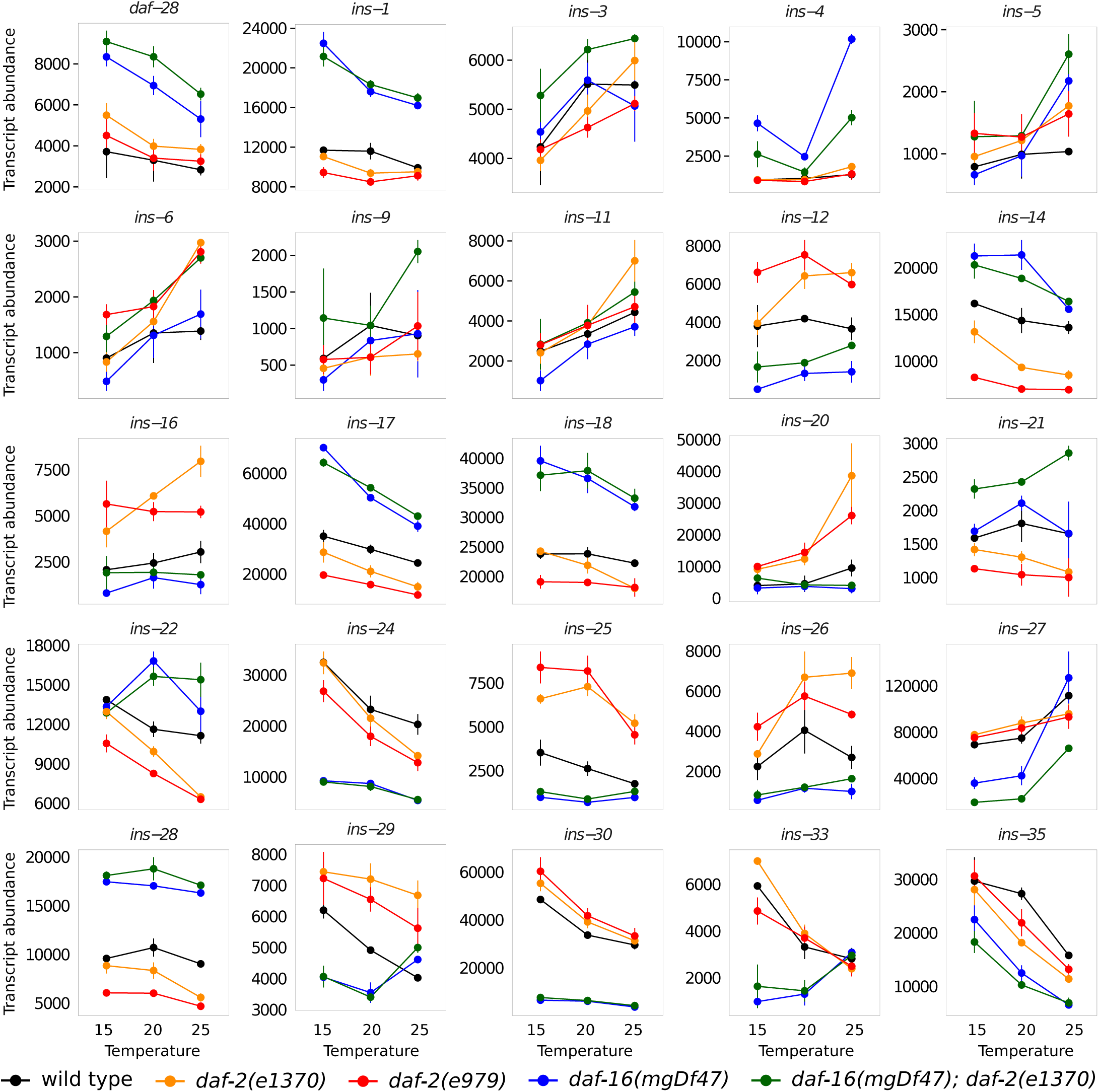
Insulin-like signaling and temperature regulate expression of the majority of insulin-like peptides during L1 starvation. Transcript abundance (arbitrary units) is plotted during L1 starvation at three different temperatures in five different genotypes with various levels of insulin-like signaling activity. Of 28 reliably detected insulin-like genes, 25 were significantly affected by *daf-16* and are plotted. 21 were significantly affected by temperature (all but *ins-2, 9, 12, 16, 21, 29, 34*). *ins-10* was affected by temperature but not *daf-16* and is not plotted here (see Tables 1, S1 and supp. data). Error bars reflect the SEM of two or three biological replicates.

Supp Data File 1. Complete normalized expression data set. Raw data is available upon request.

Supp Data File 2. Complete statistics for regulation of all genes during starvation and recovery by *daf-16* and temperature.

**Table S1.**
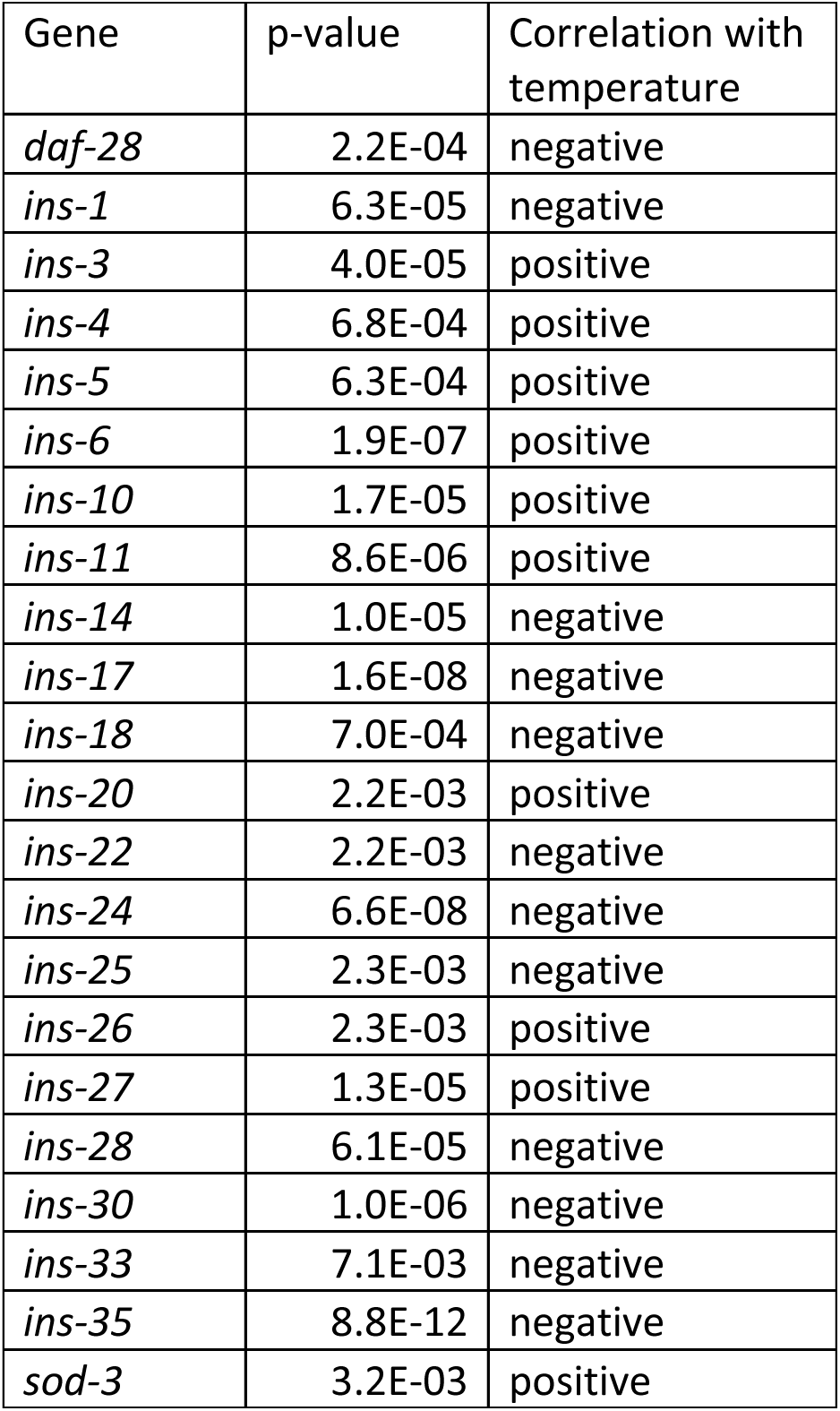
Summary of insulin-like peptides whose expression is affected by temperature during L1 starvation. The gene, the nominal p-value for the effect of temperature after controlling for genotype (not corrected for multiple testing), and whether the gene’s expression is positively or negatively correlated with temperature is presented. Only those genes with a p-value below 0.05 after correction for multiple testing are included.

